# Feedforward inhibition allows input summation to vary in recurrent cortical networks

**DOI:** 10.1101/109736

**Authors:** Mark H. Histed

**Affiliations:** National Institute of Mental Health, National Institutes of Health, 35 Convent Dr. 35/3A-203, Bethesda, MD 20892..

## Abstract

Brain computations depend on how neurons transform inputs to spike outputs. Here, to understand input-output transformations in cortical networks, we recorded spiking responses from visual cortex (V1) of awake mice of either sex while pairing sensory stimuli with optogenetic perturbation of excitatory and parvalbumin-positive inhibitory neurons. We found V1 neurons’ average responses were primarily additive (linear). We used a recurrent cortical network model to determine if these data, as well as past observations of nonlinearity, could be described by a common circuit architecture. The model showed cortical input-output transformations can be changed from linear to sublinear with moderate (∼20%) strengthening of connections between inhibitory neurons, but this change depends on the presence of feedforward inhibition. Thus, feedforward inhibition, a common feature of cortical circuitry, enables networks to flexibly change their spiking responses via changes in recurrent connectivity.

**Significance statement:** Brains are made up of neural networks that process information by receiving input activity and transforming those inputs into output activity. We use optogenetic manipulations in awake mice to expose how a transformation in a cortical network depends on internal network activity. Combining numerical simulations with our observations uncovers that transformation depend critically on feedforward inhibition – the fact that inputs to the cortex often make strong connections on both excitatory and inhibitory neurons.

## Introduction

Neurons in the cerebral cortex receive thousands of synaptic inputs and transform those inputs into spike outputs. Input-output transformations can be characterized in single cells (measuring firing rate while injecting current to produce a f-I curve, (Connors et al., 1982; Destexhe and Paré, 1999; Koike et al., 1970)), but network effects can dramatically alter input-output transformations *in vivo*. For example, ongoing network activity can create supralinearities in neurons’ input-output functions (Priebe and Ferster, 2008), strong network connectivity can create entirely linear input-output functions (Brunel, 2000; van Vreeswijk and Sompolinsky, 1996), and recurrent connections can amplify inhibition to produce sublinearity (Ahmadian et al., 2013).

In this work, we examine input-output transformations *in vivo* by first measuring spiking responses to combinations of visual and optogenetic input in the mouse visual cortex (V1). Then, to shed light on the network and circuit mechanisms of input-output transformations, we use a spiking recurrent network model. The experimental data show that excitatory neuron stimulation gives a primarily linear (additive) input-output transformation in mouse V1, which stands in contrast to sublinearity seen in monkey V1 (Nassi et al., 2015). The model shows that the cortical network can achieve both kinds of transformations with only moderate changes in local recurrent synaptic strengths. The model makes a further prediction that feedforward inhibition – input that synapses not just on excitatory but also on inhibitory neurons – allows the cortex to support both kinds of transformations.

Optogenetic stimulation can reveal how networks *in vivo* transform inputs into output. Studies using sensory stimuli alone are complicated by the fact sensory stimuli are processed by many brain regions, each of which may provide input to a cortical area under study. Combinations of sensory stimuli have, however, found that a wide range of transformations are possible, often finding evidence for normalization, a form of sublinear summation (Carandini and Heeger, 2012). A few recent studies have used direct optogenetic input to study input-output transformations, and studies in different species have observed both normalization (Nassi et al., 2015; Sato et al., 2014) and more linear summation (Huang et al., 2014), pointing to the need to understand what features of cortical networks can change input-output transformations.

Models and theoretical approaches complement experimental studies of input-output transformations, because is difficult to control connectivity in an *in vivo* cortical network experimentally. Rate-based models (Ahmadian et al., 2013; Rubin et al., 2015) have characterized the range of behaviors cortical networks can support. But not all the effects seen in rate-based models may occur in biological networks, as spiking neurons have biophysical properties that can impact input-output transformations, such as refractory periods and nonlinearities due to spike threshold. Analysis of networks of spiking neurons is most advanced for models that approximate neuronal inputs as currents and not conductances (e.g. Brunel, 2000), but input-output relationships can be modified by the changes in effective synaptic strength and Vm variability (Richardson, 2004, 2007) that occur in realistic conductance-based neurons. Therefore, we use numerical simulations of models of conductance-based spiking neurons to determine which connectivity properties might create the input-output transformations seen in our data and in past data.

Below, we first describe the experimental results from excitatory optogenetic perturbations in mouse visual cortex (Figs. 1-2), showing near-linear responses across a wide range of firing rates and visual contrast. We then describe results from the model, showing that feedforward inhibition can produce sublinearity (Fig. 3), and that with feedforward inhibition, local connectivity can allow networks to be either linear or sublinear (Figs. 4-5). Finally, we construct a model network (Fig. 6) that fits the observations, and show it is consistent with data from optogenetic perturbations of inhibitory neurons (Fig. 7). The observations are together best described by a model with feedforward inhibition.

**Figure 1:**
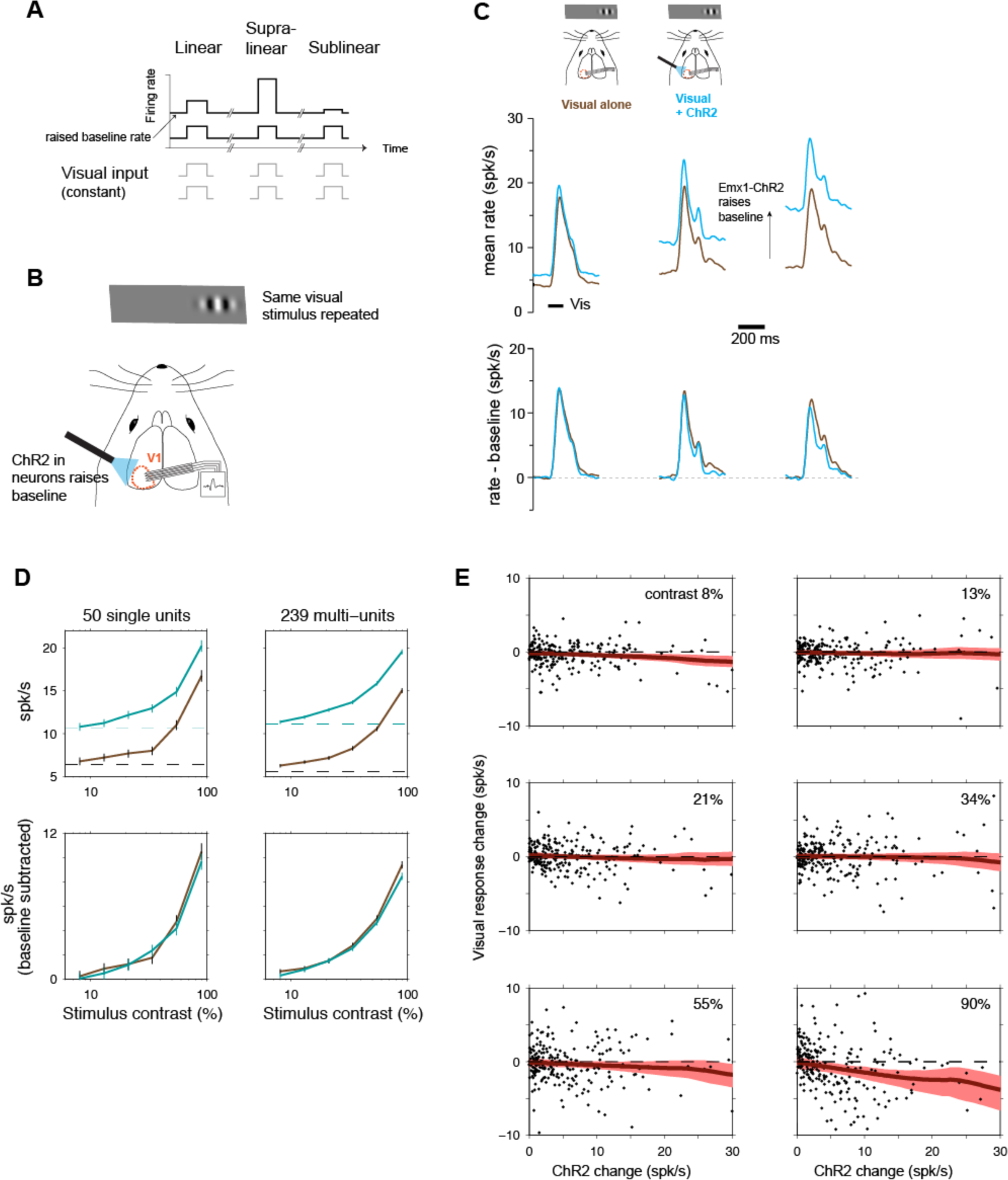
Near-linear scaling with excitatory optogenetic stimulation in mouse V1. **A.** Schematic of experimental stimulus protocol. If scaling is linear, the same input pulse produces the same response when baseline (spontaneous) rate is changed. **B.** We raise baseline rates using ChR2 in excitatory (E) neurons (Cre-dependent virus in Emx1-Cre mouse line.) **C,** Population histograms showing responses to combined ChR2 and visual (90% contrast) stimuli. Top row: columns show three groups of neurons, divided based on size of ChR2 baseline firing rate changes, left: smallest ChR2 effects (N=94; 36 single, 58 multi-units), middle: intermediate ChR2 effects (N=101; 31 single-, 70 multi-units), right: largest ChR2 effects (N=94; 28 single-, 66 multi-units), Brown: responses to visual stimulus with no optogenetic stimulus. Cyan: responses to visual stimulus when baseline rates are changed by sustained optogenetic stimulus. Bottom row: Same data as top row, with spontaneous firing rates subtracted. Visual responses differ somewhat between columns because each column is a different group of neurons, but within each group there is little response change as spontaneous rate varies. **D,** Linear scaling is seen across a wide contrast range. Top row: responses without baseline subtraction. Bottom row: baseline subtracted. Errorbars: SEM of pooled unit responses. **E,** Linear scaling is seen on average, across neurons with a variety of ChR2-induced baseline rate changes, with some weak sublinearity at the highest rate changes and highest contrasts. Y axes: difference in visual responses (relative to baseline) with and without ChR2 stimulation; dashed line at zero shows a perfectly linear response. Red: lowess regression, shaded region is a bootstrapped 95% confidence interval. Two outlier points in 90% contrast plot are omitted for visual clarity although they are included in the regression; the two outliers are shown in Fig. 2A.

**Figure 2:**
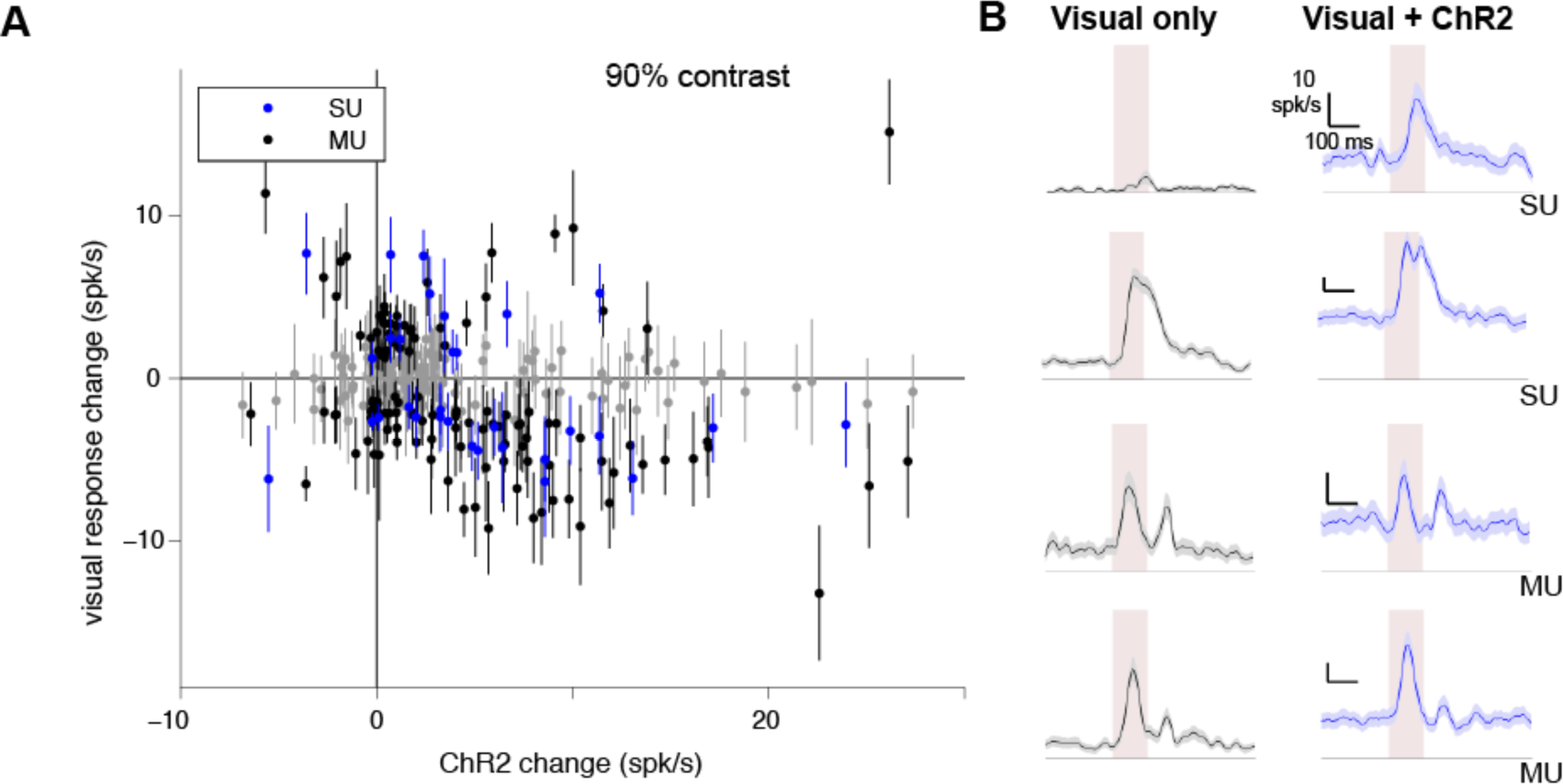
Different units can be sub- or supra-linear, though mean of population is near-linear. **A,** Unit responses to excitatory neuron optogenetic (Emx1-ChR2) stimulation, showing that many individual units are significantly supra- or sub-linear. X-axis: average firing rate change with ChR2 stimulus, Y-axis: difference between visual responses (90% contrast; each visual response measured from preceding baseline) with and without optogenetic stimulus. Errorbars: SEM. Points that are at least 1 SEM away from horizontal line at zero (linear response) are colored blue (single units; SU) or black (multi-units; MU). Points within 1 SEM of linear are gray. Data are as in Fig. 1E for 90% contrast, here with std. err. for each point, and adding on the negative Y-axis the few units that are suppressed by stimulation. 34% of single units are significantly non-linear (17/50, *p*<0.01, KS test), and 28% of multi-units are significantly non-linear (67/239, *p*<0.01, KS test). **B,** Four example units. Pink region shows visual stimulus presentation time. Shaded regions around mean response: SEM.

**Figure 3:**
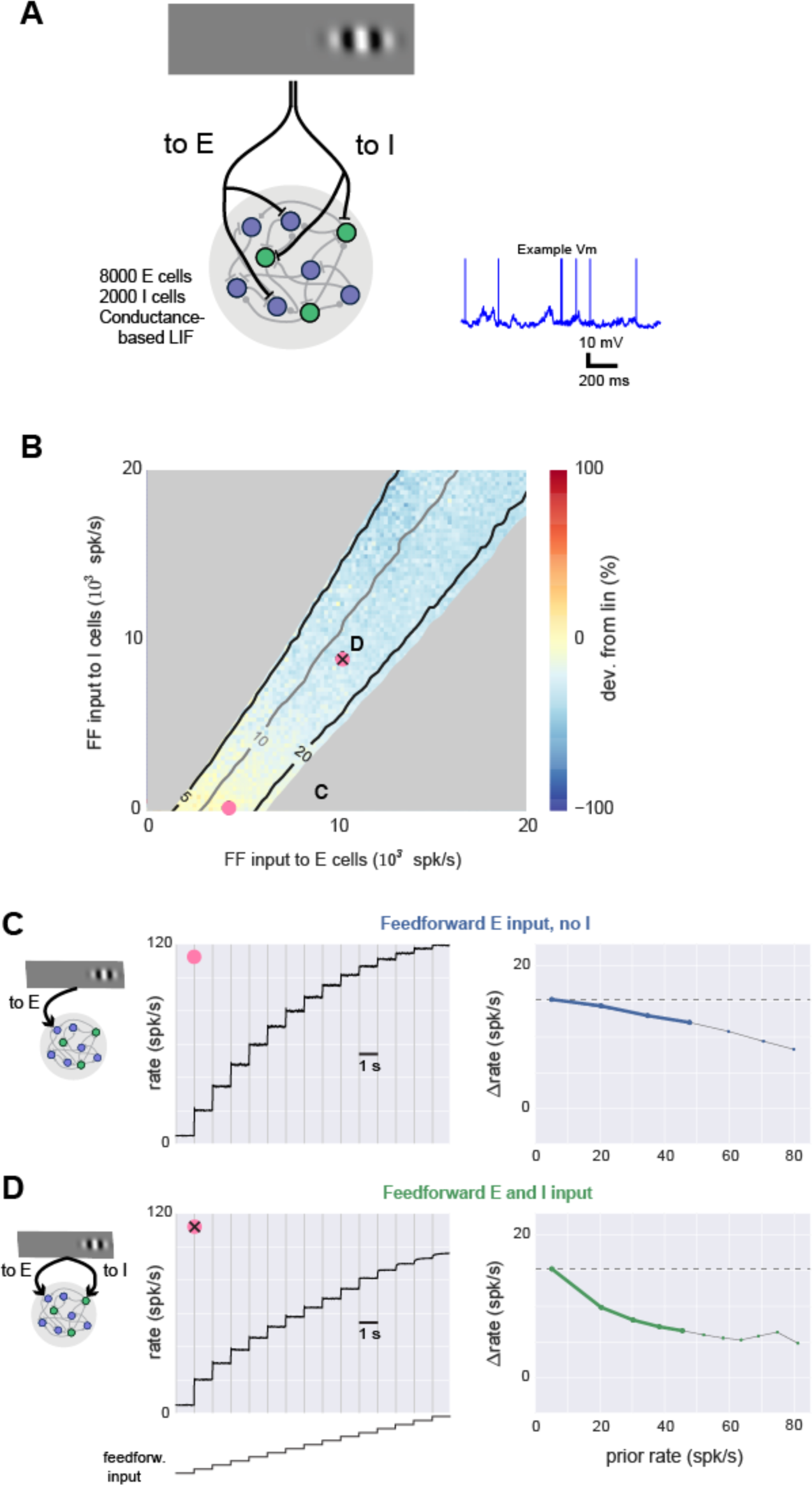
Spiking model shows sublinear scaling with feedforward inhibition. **A,** Schematic of network architecture. Blue: E cells, green: I cells. The conductance-based spiking model produces stochastic V_m_ and spikes as seen *in vivo*, and an example membrane potential (V_m_) trace from one excitatory cell is shown. **B,** Response scaling as feedforward (FF) input to E and I cells is varied. To measure response scaling, inputs to E and/or I cells with rate given by X,Y axes are delivered, and average response over all E cells is measured. Then, the E and I input rates are multiplied by a constant (here, 2) and the size of the second response is compared to the first. Percent change shown by color, yellow: second response is similar (linear), blue: second response is smaller (sublinear). Contour lines show first response (spk/s). Response rates below 5 spk/s and above 20 spk/s are masked (gray). Average spontaneous rate is adjusted to 5 spk/s (Methods), and 33% of network neurons receive external input, to approximate the sparse set of cortical neurons that typically respond to sensory inputs (Fig. 1). Pink points show E and I rate combinations used in C,D. **C,** Near-linear responses to a range of input sizes when feedforward input is provided to E cells only. Parameters here are indicated by pink dot in B, and first two responses here are the same two responses used to compute percent change shown in color there. Left panel: average rates, right panel: same data replotted showing change (spk/s) in response (y-axis) as a function of prior response (x-axis). For these plots, a linear response is a horizontal line (dashed gray line). Heavy lines: prior rates less than 50 spk/s, highlighting for visual clarity rates far from potential saturation caused by absolute refractory period (3 ms). **D,** Sublinear responses to a range of input sizes when input provided to both E and I cells. Same conventions as C. In this case, heavy green line in right panel lies farther below horizontal than heavy blue line in C, showing more sublinear scaling.

**Figure 4:**
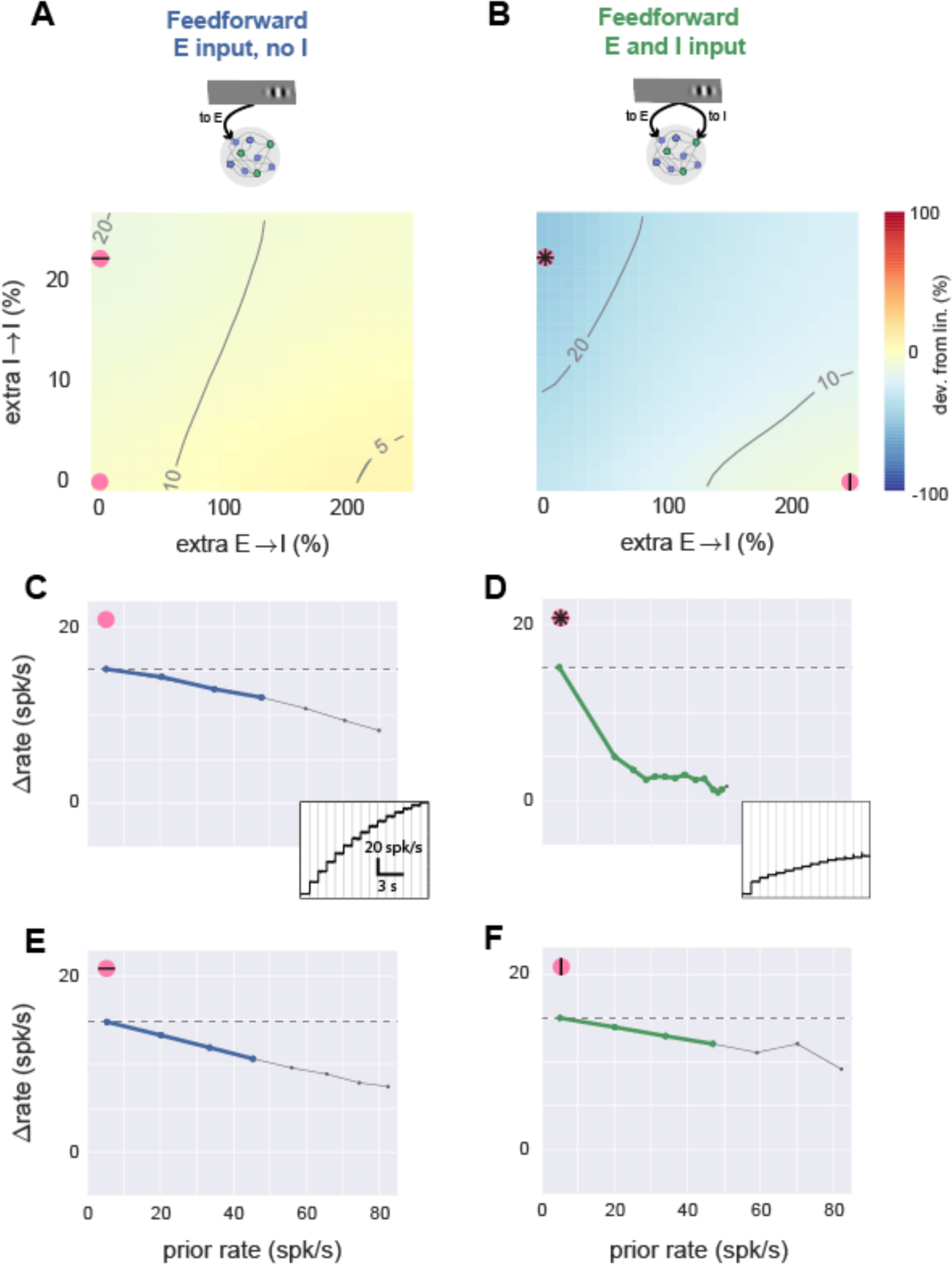
With feedforward inhibition, network model can produce linear or sublinear responses. **A,** Simulations with feedforward input to E cells only, while local network connectivity is varied. X-axis: E to I connection strength, y-axis, I to I connection strength. Axes give percent change in total synaptic input that a single cell receives from one (E or I) population (see Methods), where zero is a balanced network (e.g. Fig. 3) with equal probability of synapses onto E and I cells. Other conventions as in Fig. 3B (contour lines show evoked response to first stimulus, color shows percent difference in response to doubled external stimulus). Spontaneous rate and external stimulus rates are constant for entire panel. **B,** Simulations with feedforward input to E and I cells while local connectivity is varied. Pink symbols show parameter regions where scaling is sublinear (stronger I->I connectivity) or linear (stronger E->I connectivity). **C,** Scaling plot (response size as a function of previous rate) for parameters shown by pink dot in A: no extra local connections, feedforward E only, same parameters as Fig. 3C. Inset: timecourse of responses to the step stimulus; subtracting each rate from rate at the previous step gives y-axis in main panel. **D-F,** same plots, using parameters shown by corresponding pink dots in B. Comparing D and E shows that large sublinearity can be produced by extra I->I connections only with feedforward inhibition. Comparing D and F shows that linearity can also be achieved with feedforward inhibition if E->I connectivity is strengthened.

**Figure 5:**
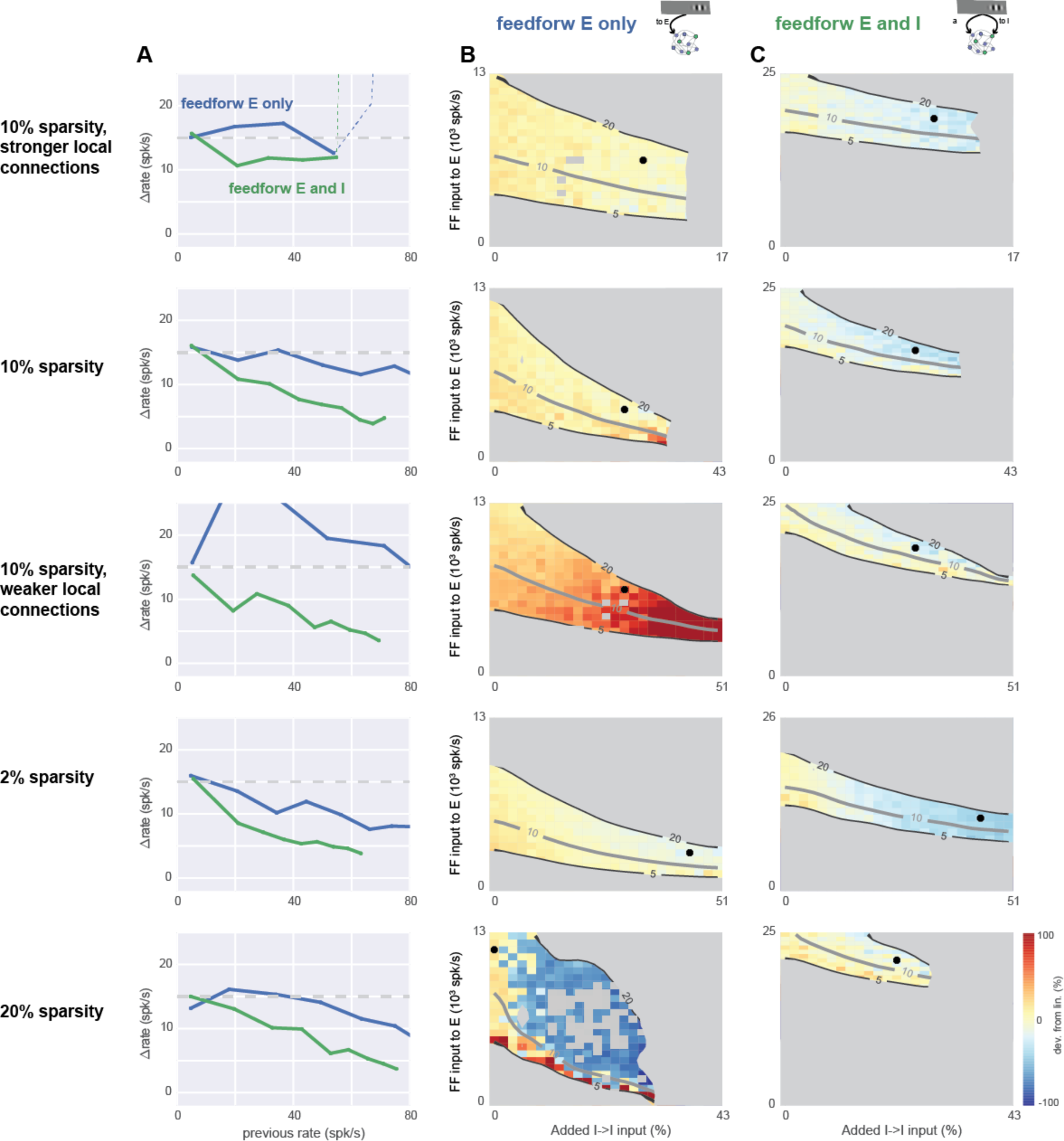
Feedforward inhibition leads to sublinearity in networks with a range of recurrent synaptic sparsities and synaptic strengths. Top row: Simulations in the conductance-based network with 10% connectivity, with strong synapses (each cell receives 10x more E and I input than in the networks of Figs. 3-4). Other rows show networks with different sparsity and synaptic strength. The network of Figs. 3-4 is the fourth row (2% sparsity, 1x strength). **A,** Scaling plots showing network response as a function of prior rate before stimulus. Blue: feedforward E input only, parameters shown in column B. Green: feedforward E and I input, corresponding parameters are shown in column C. In all rows, feedforward inhibition (green) allows more sublinearity than feedforward excitation alone (blue). Dashed line, top row: network instability (rates diverge). **B,** Average network response as I-I synaptic strength (x-axis) and feedforward E input (y-axis) are varied. No feedforward inhibition. Black dot shows parameters used to plot blue line in A (parameters chosen to maximize sublinearity). Gray regions mask areas where evoked rates are less than 5 spk/s or greater than 20 spk/s, or where network was unstable (rates diverged to maximum rate given by refractory period). Other conventions as in Fig. 3B, 4AB. **C,** network response as a function of I->I and feedforward E input, in the presence of feedforward inhibition. Individual gray squares seen in fifth row (20% sparsity) column B, inside the 5-20 spk/s contours indicate strongly irregular (non-monotonic) response scaling: strong sublinearity for at least one stimulus step, when both previous and later responses were linear or supralinear. Feedforward inhibition arrival rate to stimulated cells for each row, from top: 14k, 14k, 19k, 11k, 17k spk/s, chosen to give a 15 spk/s response for 3x the feedforward excitatory rate that alone produces a 15 spk/s response (see Fig. 3B). Fourth row (2% sparsity, same network as Figs. 3-4) uses 40% extra I->E connections to show linear responses are robust to many forms of connectivity variation.

**Figure 6:**
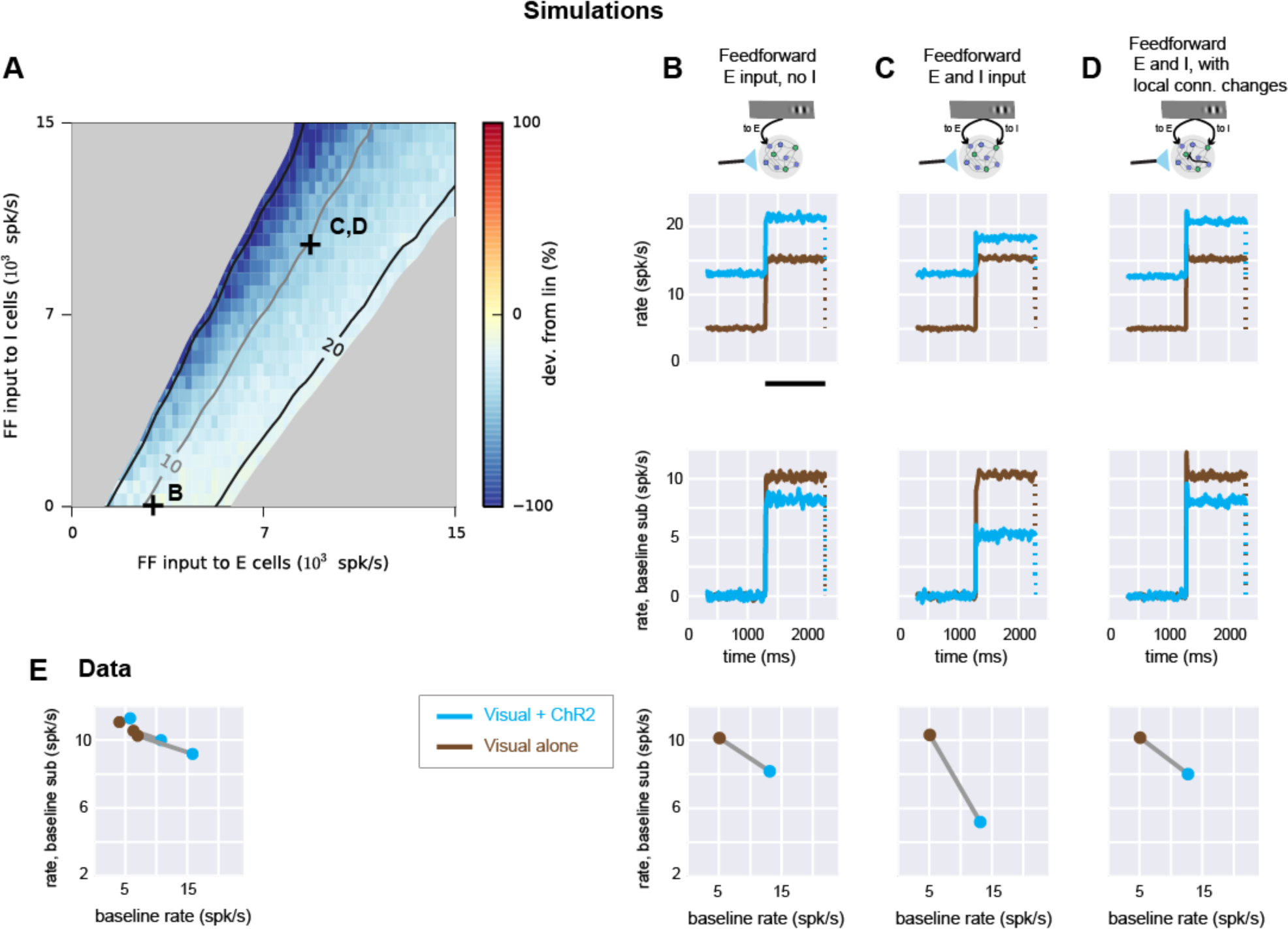
Experimental linear scaling can be replicated in networks receiving feedforward inhibition. **A,** Simulation where conductance steps (ChR2 input) and feedforward Poisson trains (visual input) are combined. Strength of feedforward E input (x-axis) and feedforward I input (y-axis) are varied while spontaneous rate is set to 5 spk/s. Connection sparsity is 2%. Other conventions as in Fig. 3B. Symbols (+) show values of E,I input used in panels B-D. **B,** network responses when feedforward input is supplied to E cells only. Top row: network responses (mean of E cell rates). Brown: feedforward Poisson (visual) input only. Cyan: conductance (ChR2) input combined with visual input. Conductance increase lasts for the full duration of the cyan trace. Visual input duration is shown by black bar (bottom of plot). Dotted line indicates rates return to previous baseline when feedforward input ends. Second row: same data as top row, with baseline rate subtracted. Third row: response (y-axis) as a function of rate before feedforward input begins (x-axis). **C,** same network simulations with feedforward input to both E and I cells (parameters marked by C in panel A). **D,** network receiving feedforward input to both E and I cells, but with stronger local connections from E to I cells (cf. Fig. 4, with similar effect for two feedforward Poisson inputs instead of feedforward input paired with conductance step as shown here). **E,** data from Fig. 1C plotted to show how responses scale as baseline is changed. Three lines (brown: no ChR2, cyan: with ChR2) are the three groups of recorded neurons shown in Fig. 1C.

**Figure 7:**
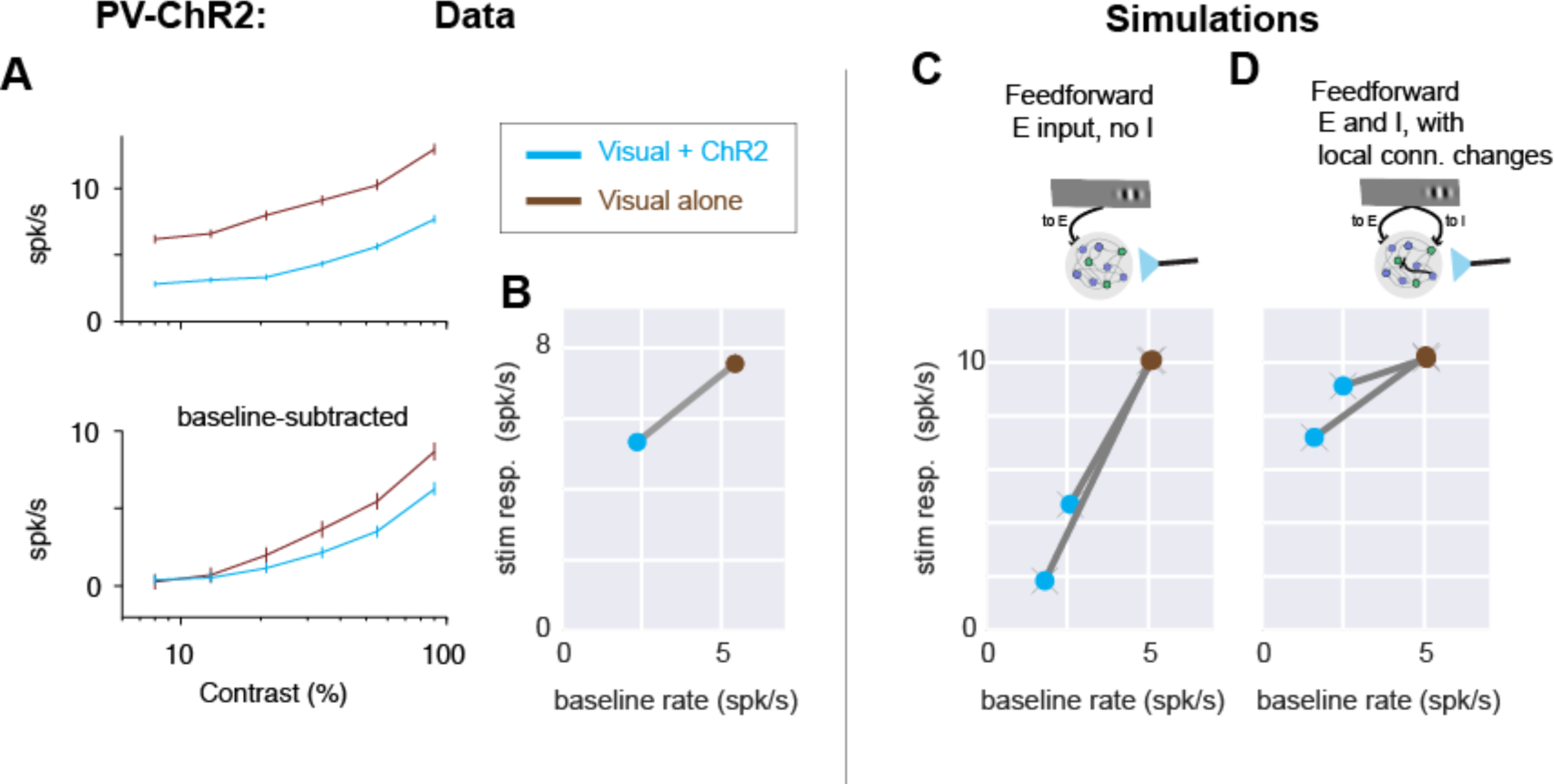
PV-ChR2 stimulation data support the recurrent model with feedforward inhibition. **A,** Moderately sublinear scaling of visual responses is seen when PV neurons are optogenetically stimulated. (Data set previously reported in Glickfeld et al., 2013). Same conventions as in Fig. 1D. N=43 units, 6 SU, 37 MU. **B,** response sizes plotted as a function of baseline rate; same conventions as bottom panels in Fig. 6B-D. Stimulation of PV inhibitory neurons lowers baseline firing rates (here 2.3X reduction), so visual + ChR2 response (blue point) is to the left of visual only (brown). **C-D,** Model (with feedforward inhibition) that best fits E neuron stimulation data also describes moderate sublinearity seen in PV-ChR2 stimulation. C, model with feedforward input to E cells only (same model as in Fig. 6B) shows very strong sublinearity. Two lines show two different strengths of optogenetic input to I cells (chosen to produce 2X or 3X decrease in baseline rates). D, model with feedforward input to E and I cells and stronger local E to I connectivity (same model as in Fig. 6D), shows a range of sublinear scaling similar to that seen in the experimental data (A-B).

## Materials and Methods

### Neurophysiology

All experimental animal procedures were conducted in accordance with NIH standards and were approved by the IACUC at Harvard Medical School. Animal breeding and surgery were performed according to the methods described previously (Glickfeld et al., 2013; Histed and Maunsell, 2013).

Neurophysiological data from Emx1-Cre animals (N=4, of both sexes but sex not recorded) were collected using the methods used in Glickfeld et al. (2013) for extracellular recordings. Briefly, animals kept on a monitored water schedule were given small drops of water (∼1 µL) every 60-120 s during recording to keep them awake and alert. The visual stimulus, a Gabor patch with spatial frequency 0.1 cycles/deg and sigma 12.5 deg, were presented for 115 ms (FWHM intensity) and successive visual stimuli were presented every 1 s. Optogenetic light pulses were delivered on alternating sets of 10 stimulus presentations (light onset 500 ms before first stimulus, offset 500ms after end of last stimulus; total light pulse duration 10.2s). A 1 s delay was added after each set of 10 stimulus presentations. Extracellular probes were 32-site silicon electrodes (Neuronexus, Inc., probe model A4x8). Recording surfaces were treated with PEDOT to lower impedance and improve recording quality. On each recording day, electrodes were introduced through the dura and left stationary for approximately 1 hour before recording to give more stable recordings. ChR2 was expressed in excitatory neurons (as described in Histed and Maunsell, 2013) using viral injections into the Emx1-Cre (Gorski et al., 2002), (Stock #5628, Jackson Laboratory, Bar Harbor, ME USA) line. Virus (0.25-1.0 µL) was injected into a cortical site whose retinotopic location was identified by imaging autofluorescence responses to small visual stimuli. Light powers used for optogenetic stimulation were 500 µW/mm^2^ on the first recording session; in later sessions dural thickening was visible and changes in firing rate were smaller, so power was increased (maximum 3 mW/mm^2^) to give mean spontaneous rate increases of approximately ∼5 spikes/s in that recording session. Optogenetic light spot diameter was 400-700µm (FWHM) as measured by imaging the delivered light on the cortical surface. Spike waveforms were sorted after the experiment using OfflineSorter (Plexon, Inc.). Single units were identified as waveform clusters that showed clear and stable separation from noise and other clusters, unimodal width distributions, and inter-spike interval histograms consistent with cortical neuron absolute and relative refractory periods. Multiunits were clusters that were distinct from noise but did not meet one or more of those criteria, and thus these multiunits likely group together a small number of single neuron waveforms.

### Experimental Design and Statistical Analysis

Spike histograms were smoothed using piecewise splines (LOWESS smoothing). To compute neurons’ visual responses (e.g. Fig. 1D, 2A), we counted spikes over a 175 ms period beginning 25 ms after stimulus onset, with a matched baseline period 175 ms long ending at stimulus onset. To test for non-linearity, for each cell we found the response count with and without optogenetic stimulation by taking the stimulus response count and subtracting the baseline count. Neurons were classified as significantly non-linear if the p-value of a two-sample two-tailed Kolmogorov-Smirnov test on the counts with and without stimulation was less than 0.01. Comparing the percent of units significant to 1% (the percentage was much higher) controls for multiple comparisons. The Emx1 dataset includes data from 100 shank penetrations (∼25 recording sessions with a 4-shank electrode). Because the inter-shank spacing was 200-400 µm, our stimuli in fixed retinotopic locations could not activate neurons on all shanks. Therefore, we included only shanks in which an average visual response > 0.2 spikes/s was measured (38/100 shanks). This gave 417 single and multi-units. We examined only units that showed a visual stimulus response (N=289; mean stimulus response-mean spontaneous > 0.2) in the absence of ChR2 stimulation. Because ChR2 expression was highest at the site of viral injection and fell off with distance, we took advantage of this variation to sort units into three groups based on the strength of local ChR2 activation (Fig. 1C). We found the average change in spontaneous rate induced by ChR2 stimulation for all units on a shank and rank-ordered the shanks. Dividing shanks into three groups based on small, medium, or large ChR2 effects yielded three nearly-equal sized groups of units receiving small, medium or large ChR2 activation. The group sizes differ by a few units because we sorted by shank, not by individual unit.

To test whether units were non-linear, we subtracted spike count around visual responses described above, subtracted

### Conductance-based spiking network model

The cortical model is a recurrent network of conductance-based leaky integrate-and-fire neurons. Example Python code and a Jupyter notebook (http://jupyter.org) are provided at [*url redacted for review; code provided on request*] that run the network simulation with all its inputs, replicating spike counts shown in Fig. 6C, bottom row. To recover the rest of the simulations in Fig. 3-7, this code can be run in parallel on a larger cluster.

Each model neuron is connected randomly to each other neuron with fixed probability (sparsity). For example, for a 10% sparsity network, each cell receives input from 10% of the excitatory cells and thus gets 0.1*8000 = 800 E inputs. Similarly, at 10% sparsity, each cell receives 0.1*2000 = 200 I inputs. As seen in the cortex, we chose the inhibitory synaptic strength to be larger than the excitatory synaptic strength. We varied both synaptic strengths and found that our conclusions are not affected by changes in E/I synaptic strength ratio. (See also Fig. 5 for effects of changing together E and I recurrent synaptic weights by an order of magnitude). We refer to this baseline set of random, sparse connections as the balancing connections. For convenience, to change local connectivity, we change the strength of a second added set of connections with the same sparsity while keeping the strength of the balancing connections constant. For example, when I->I connectivity is varied in the 2% sparsity network (e.g. Fig. 4), each I cell receives an extra 40 synapses from other I cells, and the y-axis in Fig. 4AB shows the effects of varying the weight of those 40 synapses from zero to ∼20% of the weight of the standard recurrent I->I synapses.

Each simulated neuron’s membrane potential evolves according to the following equation:

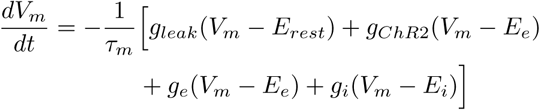

When the membrane potential *V_m_* crosses a threshold (-50 mV), a spike is recorded and *V_m_* is reset to *E_rest_* (-60 mV) for the absolute refractory period (3 ms).

Beyond the recurrent inputs from other neurons in the network (described in the model architecture above), model neurons can receive two kinds of external inputs: external feedforward inputs simulating e.g. sensory input from thalamus, and external ChR2 inputs. Feedforward (sensory) inputs are simulated as Poisson spike trains whose rates are changed by stepping to a new value, with values chosen to approximate visually-evoked changes seen in the data. ChR2 input is simulated by linearly ramping *g_ChR2_* to a new value over 2 ms, a timescale consistent with ChR2 *t_on_* (Nikolic et al., 2009), and *g_ChR2_* amplitude is varied to reproduce experimental changes in firing rate (see below). Synaptic conductances *g_e_* and *g_i_* are incremented instantaneously by a constant excitatory or inhibitory synaptic weight when a spike is fired by a recurrent or feedforward input. The conductances decay with time constants *τ_ge_* = 5 *ms* and *τ_gi_* = 10 *ms*, described by:

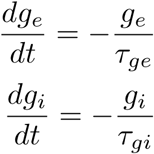

Other constants are: excitatory reversal *E_e_* = 0 mV, inhibitory reversal *E_i_* = -80 mV, membrane time constant *τ_m_* = 20 ms. Post-synaptic potential (PSP) amplitudes can vary with network activity and synaptic weight because the model neurons are conductance-based. As we varied sparsity in the network, the excitatory PSP amplitude varied over an approximately tenfold range (0.3-3.0 mV for sparsity 20% - 2%, if calculated assuming that the mean membrane potential of network neurons is -65mV.)

The sparse recurrent connections yield spontaneous activity in the network in the absence of external input (van Vreeswijk and Sompolinsky, 1998; Vogels and Abbott, 2005). To equate the spontaneous firing state of the network across different sparsity and synaptic strength, we adjust network spontaneous rate. We use an additional external Poisson excitatory input to either excitatory or inhibitory neurons to respectively raise or lower the spontaneous rate. The rate of this Poisson input is chosen via stepwise optimization to give a mean spontaneous rate across excitatory neurons of 5 spk/s. (In the 2% sparsity network, these added excitatory synapses account for only approximately 2% of the total mean conductance). For many networks, a local minimum of the parameter can be found repeatably, but for extreme values of sparsity and synaptic strength, the network is unstable and spontaneous rates are either sensitive to small perturbations or diverge. In these cases network response is not shown (e.g. gray regions, Fig. 5B-C).

Simulations were performed with the Brian package (Brette et al., 2007) on a multi-CPU cluster (the NIH HPC Biowulf cluster, http://hpc.nih.gov, or Orchestra, http://rc.hms.harvard.edu) with an integration time step of 50 µs.

## Results

### Experimental measurements in mouse V1 show linear summation

We combined visual and excitatory optogenetic input (Fig. 1A-B) by expressing channelrhodpsin-2 (ChR2) in V1 excitatory neurons using a transgenic mouse line and a Cre-dependent virus, and we used blue light pulses several seconds in duration (4-6 sec) to shift neurons’ firing rates to a new baseline. We delivered the same visual stimulus repeatedly, with and without ChR2 stimulation. We kept animals alert by giving them drops of fluid approximately once a minute, and we measured neurons’ spiking via extracellular recording with multi-site probes.

When we presented the same visual stimulus with and without optogenetic stimulation, we found that V1 neurons’ responses scaled nearly linearly (Fig. 1C) – that is, nearly the same size response was produced even as the optogenetic stimulus changed the baseline firing rate. Even for relatively large optogenetic baseline shifts (∼10 spk/s, roughly the same magnitude as the average visual response), the visual response was similar with and without ChR2 stimulation. This response implies the input-output transformation is linear (also called additive, e.g. Huang et al., 2014), meaning the sensory response produces a fixed change in firing rate above the changing baseline rate. (In contrast, if the response was sublinear, higher baseline rates would produce a smaller sensory response.) We saw nearly linear responses across a range of intensities of the visual stimulus (contrast range: 8%-90%, Fig. 1D), and we saw linear responses both in averages across single units (N=50) and multi-units (N=239). Responses became slightly sublinear in cells with the largest baseline shifts (Fig. 1E), but responses were on average within a few percent of linear (for maximum contrast, as in Fig. 1D: average sensory response changed from 10.6 spk/s to 10.2 spk/s, a -4.4% change; for single units -4.8%, for multi-units -4.1%; in contrast the average baseline rate almost doubled: 5.8 to 10.8 spk/s; change +86%).

While average neuronal responses were nearly linear, individual recorded units were often either supra- or sub-linear (Fig. 2). Units with large and small ChR2 effects are non-linear (points lie above or below the horizontal line that shows a perfectly linear response, Fig. 2A). And both SU and MU are non-linear (Fig. 2A; example timecourses in Fig 2B). With the 90% contrast visual stimulus, 34% of single units are significantly non-linear (17/50, *p*<0.01, KS test; Fig. 2A), and 28% of multi-units are significantly non-linear (67/239, *p*<0.01, KS test). Such heterogeneity in responses could arise because each neuron has slightly different local connectivity. Heterogeneity due to local recurrent connections would suggest the population average linear response is a network effect, arising from connections between excitatory and inhibitory neurons that cause them to dynamically respond to each others’ activity (van Vreeswijk and Sompolinsky, 1996). Below, using a spiking network model, we test how connectivity might lead to these observed responses.

### Other experimental work finds sublinear summation in macaque visual cortex

In contrast to this average linear scaling in mouse primary visual cortex, recent work in the monkey primary visual cortex (Nassi et al., 2015) found neural responses that were at times highly sublinear, and averages across neurons were also sublinear. (Previous work in the tree shrew and mouse has also found linearity and sublinearity, Huang et al., 2014; Sato et al., 2014). The experimental approach used by Nassi et al. does not seem to differ in important ways from our approach -- they expressed ChR2 primarily in excitatory neurons (using a CaMKII-alpha promoter strategy), stimulated an area of the cortex a few hundred microns in diameter, and they paired ChR2 and visual stimulation. Because the different results may stem from differences in cortical architecture across species, rather than differences in experimental methods, we sought to determine whether there were features of local cortical circuits that could change response scaling from linear to sublinear.

### Model network simulations identify circuit properties controlling input summation

Since it is difficult to manipulate neural connectivity *in vivo,* we used numerical simulations of conductance-based model neurons to understand how network connectivity might change response scaling. We constructed networks of 10,000 conductance-based leaky integrate-and-fire neurons, 8,000 excitatory (E) and 2,000 inhibitory (I). We chose realistic parameters for the model neurons, including sparse connectivity (initially 2%), and chose moderate synaptic strengths such that a few tens of EPSPs were required to push a neuron over threshold. (We explore a range of values of sparsity and synaptic strength below.) These sparse, randomly connected networks produce irregular and asynchronous spontaneous activity (Fig. 3A) similar to that observed experimentally (Destexhe et al., 2003; Steriade et al., 2001) and show stable responses to external inputs (Vogels and Abbott, 2005). For all simulations, we set the spontaneous average rate of the network to 5 spk/s. There are a variety of single-cell properties that could set neurons’ spontaneous rate, but we changed the spontaneous rate by supplying a small, constant amount of excitatory input that does not vary with network activity or input, to either excitatory or inhibitory neurons (see Methods).

To determine how different sorts of feedforward inputs affect neurons’ responses, we simulated external inputs to E and I cells using two input groups of Poisson spike trains whose rates could be varied independently. As expected, when we varied the external input rates, increasing input to E cells (x-axis) monotonically increased the average network response (Fig. 3B, contour lines; average of all excitatory cells in the network, a measure similar to that obtained by multi-electrode recordings) and increasing input to I cells (y-axis) monotonically decreased the average network response. However, we could hold the average response constant by adjusting the two feedforward inputs. When the average response was constant (along contour lines in Fig. 3B), we still observed changes in response scaling, and those changes depended on the amount of I input.

To assess response scaling in the model (Fig. 3), we began with a combination of E and I input that produced a 15 spk/s response (chosen because we measured experimentally an average response that peaked near 15 spk/s, Fig. 1C, D). Then, we multiplied both input rates by a single constant and measured the size of the response to the scaled input. We found that when feedforward I input is small, responses are near-linear (Fig. 3C). This is not surprising, as previous theoretical work using strong local synaptic coupling in models with binary (van Vreeswijk and Sompolinsky, 1996) or current-based neurons (Brunel, 2000) showed that networks can produce linear responses even though individual neurons in cortical networks are nonlinear (Priebe and Ferster, 2008). However, these models did not characterize the effects of varying feedforward E and I input separately, and so we varied feedforward I input in the conductance-based model. Indeed, when feedforward I input was varied, we observed deviations from linearity. Even though the spontaneous spike rate and the spike rate response to a single stimulus alone were both held constant with and without feedforward inhibition, increasing stimulus strength showed more sublinear response scaling when feedforward inhibition was present.

### Local connectivity changes summation only in the presence of feedforward inhibition

While adding feedforward inhibition induced some sublinearity, we wished to know if more dramatic nonlinearities were possible. Therefore, we next (Fig. 4) changed local recurrent connectivity between and amongst E and I populations, and measured how those connectivity changes affected response scaling. Fig. 4 shows the effects of varying two local connections (first, strength of synapses from E to I, and second, strength of synapses from I to I) to illustrate the range of effects we observed. To implement varying connectivity in the model, we added additional connections between two neuronal populations (e.g, E to I, or I to I) with the same sparsity as the network. We then varied the strength of those additional connections and measured effects on response scaling.

With only feedforward input to E cells (Fig. 4A, xC, E), we found that changing network connections did not dramatically affect response scaling. Changing the connectivity could change the gain of the network (the size of the response to a constant input, Fig. 4A, contour lines), but response scaling was nearly linear (Fig. 4A, plot is yellow throughout; Fig. 4C-D: black lines lie close to horizontal dotted line). At high firing rates, we consistently saw moderate increases in sublinearity, which seems likely to be due to effects of the 3 ms absolute refractory period. (To focus on rates well below the refractory period, we show rates above 50 spk/s as light gray lines in Fig. 4CD). We also varied all pairwise combinations of E to I connectivity, as well as feedforward E and I input strength, and found that without feedforward inhibition, responses never showed substantial nonlinearity. Thus, the linear scaling we had observed in the model when delivering input to E cells only was robust to changes in local connectivity. In sum, without feedforward inhibition, scaling was approximately linear, and local connectivity changes had little effect.

Near-linear scaling was consistently seen when feedforward input arrived to E cells, but when feedforward input arrived to both E and I cells, responses could be either linear or sublinear. When we increased local I to I connection strength (Fig 4B, y-axis), sublinearity was observed (Fig. 4D; plot parameters correspond to pink asterisk in Fig. 4B, in blue region of plot). But increased E to I connection strength (Fig. 4B, x-axis) led to increasingly linear scaling (Fig. 4E; plot parameters correspond to pink ‘|’ symbol in Fig. 4B). The sublinear scaling produced by stronger I to I connectivity was dramatic. As with all the timecourse plots (Fig. 4C-F), we chose input strength so the first firing rate response was 15 spk/s, but when I to I connectivity was increased, subsequent firing rate responses fell as low as 1 spk/s (Fig. 4D). The mechanism by which increased I to I coupling produces increased sublinearity is not yet understood. Such unintuitive changes might possibly arise from network-level effects, similar to the way E-I tracking may cause inhibitory neurons to actually decrease their activity when inhibitory neurons are excited by stimulation (Ahmadian et al., 2013), or might arise from cell-autonomous changes in conductance that leads to shunting in individual cells (Chance et al., 2002; Richardson, 2004). Further theoretical work will be required to understand why increased I-I coupling leads to increased sublinearity in spiking networks. However, it is likely that I-I connectivity changes can be achieved in cortical inhibitory neurons, as inhibitory cells modify their dendritic structure over time (Chen et al., 2011). In sum, the numerical simulations show that local connectivity changes can dramatically affect response scaling, but only in the presence of feedforward I input.

### Connectivity effects on summation do not depend on connection sparsity or strength

We next examined whether synaptic strength and connection sparsity can change the role of feedforward inhibition in response scaling. We expected that varying the total recurrent input that neurons receive would change non-linearity of responses (as predicted by theory, Ahmadian et al., 2013; van Vreeswijk and Sompolinsky, 1996), as long as the network remained stable. Therefore, we varied total input in two ways, by varying connection sparsity and by varying synaptic strength (Fig. 5). Experimental estimates of local connection sparsity range as high as 10-20% (*i.e.* each neuron connects to 10-20% of nearby neurons, Braitenberg and Schüz, 2001; Lefort et al., 2009). But the effective sparsity of connections might be lower, as connection probability in cortical networks is known to fall off with distance. Average network connection probabilities might thus be lower than the measurements, which were obtained for nearby neurons. Therefore, to examine the effects of changing connection probability, we varied sparsity between 2-20%. We found that in all these cases, adding feedforward inhibitory drive allowed more sublinear responses (Fig. 5; green lines always lie below blue lines in Fig. 5A). We observed more linear scaling when we increased the strength of all synapses together, and a bigger range of possible scaling (from supralinear to sublinear) when we decreased synaptic strength. These results show that, in networks that use a range of connection strength and sparsity, feedforward inhibition enables local E and I connectivity to have similar effects on response scaling, though the networks became more linear as connectivity strength increased.

### Summation in our data and past data can be explained by a model with feedforward inhibition

Next, we asked whether a model that incorporates realistic optogenetic input shows the same scaling dependence on feedforward inhibition we have observed. Up to this point, we have examined the behavior of simulated networks only by scaling a feedforward input (Figs. 3-5). We have implemented this feedforward input to simulate the way input spikes change conductance in neurons, by modulating the firing rate of a (Poisson) stochastic point process. Using these input spike trains, the sum of feedforward synaptic inputs in a given network neuron has substantial fluctuations about its mean. In contrast, experimental ChR2 stimulation activates many channels, and produces conductance changes with much smaller fluctuation about the mean. Thus, it might be possible that the scaling behavior we studied experimentally, with ChR2 combined with visual stimuli, would differ from the combinations of feedforward input we simulated in Figs. 3-5. To determine if there was a difference, we simulated ChR2 input by changing conductance and combined this with feedforward input (Fig. 6), and found that combinations of ChR2 and visual inputs produced qualitatively similar effects to the effects we had previously seen. Combinations of simulated ChR2 and visual input (Fig. 6A) showed slightly increased sublinearity when compared to a single scaled visual input (cf. Fig. 3B). (We also saw some slight sublinearity in our measurements of responses to combined ChR2 and visual input in mouse V1, Fig. 1) However, as with simulated visual input (Figs. 3-5), we found that with paired conductance (ChR2) and spiking (visual) inputs, more sublinearity is possible when the feedforward input combines inhibitory and excitatory targets than when feedforward input targets only excitatory neurons (Fig. 6B-C). And, in the presence of feedforward inhibition, moderate changes in network connectivity can modify scaling behavior (Fig. 6D). In sum, in the models that simulate visual input alone (Figs. 3-5), and the models that simulate combined visual and ChR2 (conductance) inputs (Fig. 6), the role of feedforward inhibition and I-I connectivity in response scaling is similar.

We next asked what combinations of connectivity and feedforward input could describe both our data and past measurements. We constructed a model with combined visual (spiking) and ChR2 (conductance) inputs, and fit evoked rates to our data. Our data (Fig. 6E) was well-matched by the simulations that showed small sublinearity (Fig. 6A-D). The data was similar to two different sets of network simulation parameters. Networks with only feedforward excitation showed responses that paralleled the data (Fig. 6B). But networks with both feedforward excitation and inhibition could also describe our data when the network local connectivity was adjusted (Fig. 6D). Since feedforward inhibition is a common feature of cortical networks in many species (Douglas and Martin, 2004), a model using feedforward inhibition seems a good choice to describe experimentally measured response scaling. Further, with feedforward inhibition, changes in local (e.g. I-I) connectivity can change response scaling from linear to sublinear, describing not just our data but also past data. These simulations show that a wide regime of cortical scaling behavior, from linear (as seen here in mouse V1 and also in the tree shrew (Huang et al., 2014)), to strongly sublinear (as seen in primate V1, Nassi et al., 2015), can be achieved by a model with feedforward inhibition. In sum, the simulations show that a model with feedforward inhibition can describe both our data and past observations.

### PV neuron stimulation effects are explained by the model with feedforward inhibition

We next tested the model against data obtained by pairing visual and optogenetic stimulation of parvalbumin-positive (PV) cells. A majority of cortical PV inhibitory neurons are soma-targeting fast-spiking basket cells (Kawaguchi and Kubota, 1997; Tremblay et al., 2016), which are well-positioned to act as the balancing population in the network models. We found that stimulating PV neurons with ChR2 in awake mice produces a moderate suppression of visual responses, with a larger change in baseline rates than in stimulus responses. As before, we measure the visual response relative to the preceding baseline firing rate, which is changed by optogenetic stimulation. The optogenetic stimulation lowered the baseline firing rate by a substantial amount (from 5.4 spk/s to 2.4 spk/s, a 57% reduction), and reduced the response to a high contrast visual stimulus by a smaller amount (from 7.6 to 5.3 spk/s or 29%; Fig. 7A-B).

We then used this PV-ChR2 stimulation data to determine which models in Fig. 6 fit both the excitatory and PV stimulation mouse V1 data. As with the simulations in which excitatory neurons received ChR2 (conductance) input, we simulated the effects of optogenetic stimulation of PV cells by delivering a conductance input to PV neurons in the models. We adjusted the size of the conductance input to match the firing rate changes we saw in the data (Fig. 7C). The two models that fit the near-linear responses to excitatory stimulation (Fig. 6) are the model without feedforward inhibition (Fig. 6B), and the model with feedforward inhibition and local synapses adjusted to produce near-linear responses (Fig. 6D). For each of those two models, we simulated optogenetic input to PV cells, and measured the change in visual response size with and without optogenetic PV input. We found that the model without feedforward inhibition disagreed with the PV-ChR2 data, displaying very strong suppression (Fig. 7C). Only the model with feedforward inhibition (Fig. 7D) showed the same scaling (moderate suppression) seen in the PV-ChR2 data. The reduced suppression in the model with feedforward inhibition might be due to a smaller proportion of PV total input coming from optogenetic stimulation in that model, compared to the model where PV cells receive no direct feedforward input. In sum, optogenetic perturbations of excitatory and PV-positive cells are described by a cortical recurrent network model that requires feedforward inhibition.

In sum, our data shows that average response summation for excitatory input in mouse V1 is close to linear, even though individual cells can be nonlinear. Linear summation holds even for substantial shifts in firing rate (ChR2-induced firing rate changes of 10-15 spk/s, approximately the same size as the maximum visual response, Fig. 1). Using a numerical model of conductance-based spiking neurons, we find that response scaling is affected dramatically by synaptic connectivity. Moderate changes in synaptic coupling (∼20%) between inhibitory cells can change response scaling from linear to sublinear (Figs. 4-6). Further, the change in inhibitory-to-inhibitory (I-I) connectivity that leads to sublinear summation only yields such sublinear summation in the presence of feedforward inhibition.

## Discussion

It might seem surprising that we observed linear responses and not divisive normalization, where adding an additional stimulus yields reduction of the responses to a single stimulus. This form of sublinear summation has been observed in different visual cortical areas of several species. Linear summation, on the other hand, is also commonly seen at various stages of sensory systems, and both linear and sublinear responses may be useful at different levels (Carandini and Heeger, 2012). Linear summation may be more desirable when responses at different locations should receive equal weight, as when an organism must sensitively detect a distant predator, or when spikes that occur at different times should produce the same downstream effect. In fact, computer vision systems often use both linear and normalization steps in distinct layers or networks (Carandini and Heeger, 2012; Yamins and DiCarlo, 2016). Experimentally, normalization is usually measured with sensory stimuli, not with direct cortical input, and thus normalization might partially depend on subcortical (e.g. thalamic gain control, Bonin et al., 2006) or feedback effects.

In fact, the linear responses we observed with excitatory optogenetic stimulation in mouse primary visual cortex are similar to those seen in tree shrew visual cortex (Huang et al., 2014), but are different than the sublinear responses seen in macaque visual cortex (Nassi et al., 2015). Our simulations show that a broadly similar cortical architecture can support both kinds of scaling of feedforward input, subject to moderate adjustments in local connectivity. The linear responses we saw in the mouse differ from those of Sato et al. (2014), who also delivered combinations of excitatory optogenetic and visual input to mouse V1 neurons and found sublinearity under certain conditions. However, Sato et al. used an experimental approach different than the other three studies (macaque, tree shrew and our study in mouse), in which they optogenetically elicited antidromic input spikes by stimulating the contralateral hemisphere from which they were recording. Comparing these two types of input may shed additional light on how cortical circuits transform inputs to outputs.

To stimulate many V1 neurons, we delivered optogenetic input to multiple neurons simultaneously. We used a blue light spot a few hundred µm in diameter, comparable to the region of mouse V1 activated by our small visual stimulus. Many neurons in the cortex change their firing rate in response to even small sensory stimuli (Bonin et al., 2011; Van Essen et al., 1984). Anatomically, sensory input that arrives to multiple cells is common, as in the case of divergent feedforward thalamic input to the cortex (Reid, 2001). Single axons from the thalamus often ramify across several hundred microns of the cortex (Braitenberg and Schüz, 2001; Garraghty and Sur, 1990), and thalamic axons projecting to the visual cortex can make synapses on dozens of excitatory cortical cells (Freund et al., 1989).

Optogenetic stimuli may lead to firing rate changes in other parts of the brain besides the area stimulated. But perhaps because the majority of synapses made by cortical neurons are within the same cortical area, local intracortical effects for optogenetic stimuli like these have been observed to be larger than effects on the visual thalamus (Li et al., 2013; Olsen et al., 2012). This is true even though the visual thalamus (dorsal lateral geniculate) receives a large proportion of all projections out of V1 (Reid, 2001). Thus, the neurons best suited to act as the recurrent population in the model may be other V1 neurons, and perhaps even neurons within a few hundred microns of the neurons receiving input, where the probability of recurrent connectivity is highest (Lefort et al., 2009). However, other neurons in the brain could also in principle contribute to the recurrent population.

Our results show that network mechanisms can contribute to response summation. The model neurons are leaky integrate-and-fire neurons, so individual model neurons sum their subthreshold inputs linearly, and the nonlinear spiking responses we characterize likely arise from how E and I neurons interact. We chose this model architecture because we judged it the simplest model that could capture both excitatory-inhibitory interactions and also single-cell nonlinearities due to refractory period, Vm fluctuations, spike threshold, and conductance changes (Chance et al., 2002; Richardson, 2004). There are, however, other single-cell mechanisms, such as short-term synaptic plasticity or dendritic nonlinearity (Häusser et al., 2000; Silver, 2010) that might additionally contribute to even more nonlinear summation, both below threshold and in spike responses. On the other hand, dendritic nonlinearities might also have roles that do not affect scaling, as for example nonlinearities can be used to amplify distant input synapses so that different synapses produce equal responses at the soma (Katz et al., 2009).

We adjusted synaptic coupling between (E and/or I) populations by changing the strength of a set of fixed connections between the desired populations. Because in sparse networks like this neurons share only a small fraction of their input, we expected increases in synaptic strength to achieve the same qualitative result as adding new synapses, even if the two types of changes may not have exactly proportional effects on the behavior of the network. Fig. 5 shows that feedforward inhibition allows more sublinearity across changes in both synaptic strength and synapse number.

Feedforward inhibition is included in the canonical cortical microcircuit framework (Douglas and Martin, 2004) because it is a stereotypical feature of many cortical areas. In sensory cortical areas, including the visual cortex, it has been observed that input thalamic neurons make synapses both onto excitatory principal cells and onto inhibitory basket cells. Such feedforward inhibitory connectivity has been observed both with anatomical and physiological methods (Isaacson and Scanziani, 2011). Since inhibitory basket cells project strongly back to excitatory cells, inhibitory changes due to thalamic input arrive to principal cells a few milliseconds after the first excitatory changes. This delay of a few milliseconds between the arrival of excitation and inhibition can be used to align spike outputs of cortical neurons (Cruikshank et al., 2007; Gabernet et al., 2005; Swadlow, 2003; Tiesinga et al., 2008). Beyond shaping the timing of spike responses, however, it has been previously noted that feedforward inhibition might also be used to control the magnitude of spiking responses to thalamic input. Douglas et al. (Douglas et al., 1995) proposed that spike responses can be shaped by preferential amplication of either excitation or inhibition in cortical recurrent networks, where amplification might arise by connections within populations of excitatory or inhibitory neurons. Ahmadian and Miller (2013) later showed that rate-based networks with an excitatory and inhibitory term that are stable (so that the network does not e.g. diverge and become epileptic) have regimes of both linearity and sublinearity, although it is not yet clear which of these regimes spiking networks operate in, and which cellular or synaptic parameters affect summation. In Ahmadian and Miller’s model, individual cells can be supralinear (Priebe and Ferster, 2008), but when external drive arrives to multiple cells, supralinearity is also seen when recurrent connections are weak and thus excitation and inhibition are not strongly coupled. This may explain why we saw supralinear responses only in the model network with the weakest synaptic connectivity (Fig. 5).

Substantial recurrent intracortical response is elicited by sensory input, with approximately 2/3^rd^ of synaptic input after a sensory stimulus arising from recurrent synapses (Li et al., 2013; Lien and Scanziani, 2013). If recurrent connectivity is very strong, previous modeling results (Renart et al., 2010; van Vreeswijk and Sompolinsky, 1996) predict that excitatory and inhibitory populations are forced by the strong coupling to track each others’ activity closely, resulting in linear responses. In accord with this prediction about strongly-coupled networks, we observed increasing linearity when we increased synaptic strength (Fig. 5) as long as the network remained stable. However, for very strong recurrent connectivity, feedforward connectivity must also be very strong to drive any response (Ahmadian et al., 2013; see also our Fig. 5), which appears non-physiological (Li et al., 2013; Lien and Scanziani, 2013). Our simulations use synapses of moderate size (order 1mV with 2% sparsity as in Figs. 3,4,6 and Fig. 5 row 4; see Methods), requiring tens of PSPs to combine to produce a spike, as seen in cortical neurons (Barral and Reyes, 2016). These observations suggest that the differences in scaling we observed occur in a range of moderate synaptic strengths: low enough to avoid obligate linearity, and high enough to allow recurrent connections to contribute substantially to network input-output functions.

We found that a network model can link local connectivity to network physiological responses in ways that might be difficult to predict without the model. It has been difficult to measure many of the synapses in a brain volume, but connectomic methods (Briggman et al., 2011; Lee et al., 2016) promise to make such comprehensive synaptic mapping possible even in column-sized volumes of the cortex. Combining approaches for controlling input with methods to measure connectivity will be useful to shed light on an important part of brain computation – the input-output transformations of populations of connected cells.

## Acknowledgements

We thank Nicolas Brunel and Alessandro Sanzeni for discussion and comments, Lindsey Glickfeld, Alex Handler, and Oliver Mithoefer for assistance with neurophysiology, John Maunsell for support, and Bruno Averbeck, Barry Richmond, Lex Kravitz, and Carson Chow for comments on the manuscript. This work was funded in part by the Intramural Research Program of the NIMH and by U01 NS090576 (BRAIN Initiative). The authors declare no competing financial interests.

Conflicts of Interest: none

